# Reproductive fitness is associated with female chronotype in a songbird

**DOI:** 10.1101/2022.07.01.498449

**Authors:** Robyn J. Womack, Pablo Capilla-Lasheras, Ciara L. O. McGlade, Davide M. Dominoni, Barbara Helm

## Abstract

Research on biological rhythms has revealed widespread variation in diel timing within populations. Repeatable individual chronotypes have been linked to performance in humans but, in free-living species, benefits of chronotype are poorly understood. To address this gap, we investigated fitness correlates of incubation patterns in female songbirds (great tit, Parus major) at urban and forest sites. We confirm repeatable chronotypes (r ≥ 0.31) and show novel links between chronotype and reproductive success. In both habitats, females that started activity earlier in the day raised more fledglings. We also observed that forest females started their day at a similar time throughout the breeding season, whereas urban females tied their onset of activity closely to sunrise. Our study points to possible mechanisms that underlie chronotype variation and provides sought-after evidence for its relevance to fitness.

## Introduction

Due to the rotation of the Earth around its axis, no environments are completely constant across the 24 hour day. Hence, for organisms, appropriate diel timing of activities and physiology relative to environmental cycles is thought to be important for fitness (Kronfeld-Schor & Dayan, 2003). Still, inter-individual differences in diel activities can be large, whereby individuals display highly consistent temporal phenotypes called “chronotypes” (Alós et al., 2017; Helm et al., 2017; Roenneberg et al., 2003). Chronotype, first defined for laboratory rodents (Ehret, 1974), has gained major research importance in human studies, where millions of subjects have been scored (Roenneberg et al., 2019). Human chronotype has been associated with genetic variants (e.g., in clock genes), performance, and physical and mental health (Jones et al., 2019; Roenneberg et al., 2003). For example, in athletes, performance depends on chronotype and can be enhanced by modified wake-up time (Facer-Childs & Brandstaetter, 2015).

Interest in chronotype is rapidly increasing in ecology and evolution (Alós et al., 2017; Helm et al., 2017; Maury et al., 2020) fuelled by remote and automated tracking technology (e.g., transmitters or on-site loggers (Dominoni et al., 2013; Graham et al., 2017; Maury et al., 2020)). Simultaneous data collection from many individuals is paving the way for studying fitness implications of particular chronotypes, the mechanisms that generate them, and the maintenance of inter-individual variation (Hau et al., 2017; Helm et al., 2017; Martorell-Barceló et al., 2018). While ecological data are becoming increasingly available for chronotype, our understanding of the causes and consequences of its variation has been hindered by conceptual challenges, and by bias in the sex and traits investigated.

Conceptually, disentangling factors that contribute to variation in chronotype requires engaging with the complexity of diel timing. Timing is based on circadian rhythms which closely interact with ambient light (De Coursey, 2004). This ancient clock system integrates genetically controlled molecular clocks with various sensory pathways, primarily those that perceive and transduce light (Cassone et al., 2017; Helm, 2020; Stevenson & Kumar, 2017). Through further physiological pathways, additional environmental features (e.g., ambient temperature, predation risk) and state (e.g., nutrition, reproductive stage) modify timing (Helm et al., 2017). Chronotype is a phenotype defined by consistent timing of an individual’s rhythmic features (e.g., activity onset), relative to an external temporal reference and to conspecifics measured under similar conditions (Helm et al., 2017). The external reference is a fixed environmental phase in the diel cycle at the location of an animal. Choosing an external reference is, however, challenging. Chronotype in human studies usually refers to time after midnight (hereafter called ‘clock’ chronotype) (Jones et al., 2019; Roenneberg et al., 2003). This also works well for some other species that repeat diel routines at a relatively fixed time of day, for example, seabirds that under continuous polar light return to breeding colonies at roughly constant clock time (De Coursey, 2004; Huffeldt & Merkel, 2016). Yet, many species time their activity by tracking changing features of the natural light environment, such as sunrise and sunset (Bennie et al., 2014). Therefore, most ecological studies use annually and spatially variable aspects of the solar day (e.g., sunrise) as external references to calculate chronotype (hereafter called ‘relative’ chronotype) (Graham et al., 2017; Maury et al., 2020). Species and even local populations may differ in the extent to which they time their activities based on fixed (i.e., ‘clock’ chronotypes) or temporally changing features of the environment (i.e., ‘relative’ chronotype) (Da Silva et al., 2014; Da Silva & Kempenaers, 2017; Dominoni et al., 2013; Helm et al., 2017). Furthermore, animals typically adjust their behavioural responsiveness to light conditions over time, space, and life-cycle stage, so that use of relative chronotype can obscure consistent variation in timing (Daan & Aschoff, 1975). Thus, neither reference fully captures the animals’ chronotypes (Da Silva & Kempenaers, 2017; Shaw & Cresswell, 2014) and variation in both clock and relative chronotypes should be investigated in parallel to understand variation in chronotype in wild animals. Such integrative research is currently missing.

Secondly, investigation of chronotype-fitness associations must be broadened in scope. Fitness studies on wild chronotypes have until recently mostly focused on males (but see Maury et al. (2020)), partly due to extra-pair mating and to conspicuous features such as courtship, song and ornaments (Hau et al., 2017; Pagani-Núñez & Senar, 2016). For example, avian observational and experimental studies suggest that early-active males may sire more extra-pair young than late-active males (Greives et al., 2015; Kempenaers et al., 2010), and that such differences could be based on endogenous circadian clocks (Helm & Visser, 2010). However, females are disproportionately more involved in reproductive activities (Mace, 1985), and mating represents but a fraction of factors that shape the fitness landscape of chronotype. For example, offspring must develop to sexual maturity, potentially requiring extensive parental care, and adults must survive, forage and maintain sufficient body condition to generate offspring. Thus, data are needed for implications of female chronotype, and for diverse life-cycle stages.

To investigate fitness implications of chronotype, birds offer excellent study opportunities because their conspicuous behaviours and often nest-bound reproductive outcomes are often easily trackable. In this study, we leverage data from wild birds to i) disentangle factors that contribute to explain variation in clock and relative chronotype, and ii) newly document links between chronotype and reproductive success. Here, we examine a well-studied songbird whose chronotype has been shown to be repeatable, the great tit (Parus major; Graham et al., 2017; Meijdam et al., 2022; Stuber et al., 2015). We inferred female chronotype through measuring behaviour during incubation, a critically important post-zygotic stage of avian reproduction, while monitoring reproductive success (Capilla-Lasheras, 2018; Graham et al., 2017; Gwinner et al., 2018; Maury et al., 2020). Because features of the environment can influence chronotype, we included data from two habitat types, urban and forest, which often affect the diel rhythm of animals (diurnal animals in urban habitats often have earlier chronotypes; e.g., Dominoni et al., 2013; Miller, 2006). Our analyses also control for additional sources of environmental variation (e.g., temperature; Dominoni et al., 2014; Lehmann et al., 2012) and breeding conditions that are known to influence variation in chronotypes (Cooper & Voss, 2013; Gwinner et al., 2018).

The great tit is a small passerine species, widely distributed across Europe and Asia. Great tits reproduce every year, lay one clutch per year in our study area and are female-only intermittent incubators. They spend nights on their nests, but at daytime alternate between nest attendance (i.e., on-bouts) and foraging outings (i.e., off-bouts) (Diez-Méndez, Sanz, et al., 2021). From small temperature loggers inserted into nests of urban and forest-breeding great tits, we first infer both clock and relative chronotype of incubating females and assess consistency of chronotype across the breeding season. As our measure of chronotype, we focus on activity onset (time of the first incubation off-bout of the day), which in birds is particularly robust and sometimes associated with male fitness (Dominoni et al., 2013; Graham et al., 2017; Hau et al., 2017; Kempenaers et al., 2010; Pagani-Núñez & Senar, 2016), but we also report end of activity (time of the last incubation on-bout of the day) and duration of activity (difference in time between activity onset and activity end). Secondly, we link incubation chronotype to reproductive success from these same nests to test associations between female chronotype and fitness. Our research spans three years and multiple breeding locations in Scotland, ranging from oak forests to urban settings.

## Methods

### Study populations and field protocols

We studied five nest-box breeding populations of great tits (nest-box details: Woodcrete material, 260H x 170W x 180D, hole diameter 32 cm, Schwegler, Germany) during the breeding seasons of 2016, 2017 and 2018 (April to June). Three study populations were located in ancient deciduous forests, dominated by oak species (Quercus sp.): Scottish Centre for Ecology and the Natural Environment (SCENE; n = 28 nest-boxes included in the study; 56° 7’ N, 4° 36’ W), Sallochy Forest (n = 8 nest-boxes included in the study; 56° 7’ N, 4° 36’ W) and Cashel Forest (n = 31 nest-boxes included in the study; 56° 6’ N, 4° 34’ W). The remaining two populations were situated in an urban park (Kelvingrove Park; n = 14 nest-boxes included in the study; 55° 52’ N, 4° 16’ W) and a suburban park (Garscube estate; n = 9 nest-boxes included in the study; 55°54’ N, 4°19’ W) within the city of Glasgow (UK). Both urban sites contained a mixture of open land, small shrubs, and sparse woodland with introduced and native tree species. For further details on the study sites, see Branston et al., (2021) and Jarrett et al., (2020).

All nest-boxes were checked weekly from April 1 for signs of nest building activity and egg laying. Once a new completed clutch was detected, we calculated the date of clutch completion (from the number of eggs present between two consecutive nest-box visits, assuming that females laid one egg per day). After the estimated earliest possible date of hatching (assuming a minimum incubation length of 14 days from the date of clutch completion; Gosler 1993), nest-boxes were checked every other day, allowing determination of exact date of hatching based on nestling presence and appearance. Thirteen days after hatching, all nestlings within a brood were weighed (± 0.01 g) and ringed for individual identification (n = 57 broods of 13-day old nestlings). Nest-boxes were checked again > 21 days after hatching to determine the number and identity of any dead nestlings remaining in the nest. As our sample size varied slightly per each trait under investigation (see below), we provide a breakdown of sample size per habitat, year and trait in Table S1. Sunrise and sunset times at SCENE (56° 7’ 46’’ N, 4° 36’ 46’’ W) and Glasgow (55° 52’ 11’’ N, 4° 16’ 56’’ W) were obtained from www.timeanddate.com. Our data are collected from individual nest-boxes, rather than from identified females. Thus, some individuals might have been recorded in multiple years. Given that our study was spread across five sites over three years, the potential bias introduced by this methodological limitation is expected to be minimal (Table S1).

**Table 1.** Likelihood-ratio test (LRT) results and model coefficients for predictors explaining variation in clock time of female onset of activity (i.e., time of first incubation off-bout; n = 729 days of incubation). Significant terms are highlighted in bold. LRT results for ‘Habitat’, ‘Incubation start Date^1^’ and ‘Days before hatching^1^’ are not provided as these terms were part of a significant interaction present in the final model. The interactions ‘Incubation start date^2^×Habitat’ (χ^2^ = 0.98, p = 0.323) and ‘Days before hatching^2^×Habitat’ (χ^2^ df = 1 = 1.01, p = 0.316) were not significant and were df = 1 dropped from the model. Model coefficients (‘Estimate’) for clock time onset of activity (given in min after 00:00 h) are shown along with standard errors (‘SE’) and 95% confidence intervals (95% CI). Superscripts ‘^1^’ and ‘^2^’ refer to linear and quadratic terms, respectively.

| Fixed effect | Estimate | SE<sup>A</sup> | 95% CI<sup>A</sup> | $\chi^2$ | df<sup>A</sup> | p |
| --- | --- | --- | --- | --- | --- | --- |
| **Intercept** | 324.12 | 7.91 | 308.62, 339.63 |  |  |  |
| **Incubation start date<sup>2</sup>** | 3.64 | 54.35 | -102.88, 110.17 | 0.00 | 1 | 0.948 |
| **Incubation start date<sup>1</sup>** | -365.30 | 72.17 | -506.74, -223.85 |  |  |  |
| **Days before hatching<sup>2</sup>** | -44.97 | 19.83 | -83.82, -6.11 | 5.11 | 1 | **0.024** |
| **Days before hatching<sup>1</sup>** | -44.63 | 40.15 | -123.33, 34.07 |  |  |  |
| **Mean daily temperatures** | 0.48 | 0.35 | -0.20, 1.17 | 1.87 | 1 | 0.172 |
| **Clutch size** | -0.13 | 0.89 | -1.89, 1.62 | 0.02 | 1 | 0.883 |
| **Habitat** |  |  |  |  |  |  |
| *Urban* | — | — | — |  |  |  |
| *Forest* | 7.54 | 4.61 | -1.50, 16.58 |  |  |  |
| **Incubation start date<sup>1</sup>×Habitat** |  |  |  | 6.02 | 1 | **0.014** |
| *Incubation start date<sup>1</sup>×Forest* | 290.09 | 113.67 | 67.30, 512.87 |  |  |  |
| **Days before hatching<sup>1</sup>×Habitat** |  |  |  | 8.71 | 1 | **0.003** |
| *Days before hatching<sup>1</sup>×Forest* | 143.60 | 48.31 | 48.90, 238.30 |  |  |  |

<sup>A</sup> SE = Standard Error, CI = Confidence Interval, df = degrees of freedom for likelihood-ratio test.

### Incubation temperature data

To quantify incubation behaviour in female great tits, we deployed small temperature loggers (iButtons DS1922L-F5, Thermochron) inside their nests (Figure S1). We programmed iButtons to record temperature (± 0.0625 °C) every 3 min (Capilla-Lasheras, 2018). iButtons were placed carefully next to the eggs (after the third egg of the clutch had been laid), covered with a small piece of white cloth, and attached to the base of the nest by a green wire anchored by a small fishing weight (Figure S1).

### Environmental temperature data

To control for variation in environmental temperature when quantifying incubation behaviour (Capilla-Lasheras, 2018), daily mean temperatures for the breeding seasons of 2016, 2017 and 2018 were obtained from the UK Met Office for an area close to our forest sites (Tyndrum [56°25’N, 4°42’W] and city sites (Bishopton [55°54’N, 4°30’W]). We also incorporated daily mean temperatures in our statistical models explaining variation in incubation behaviour (details below).

### Quantification of incubation behaviour

Some individuals removed iButtons from the nest cup and pushed them to the side of the nest-box, so that these iButtons did not record incubation temperature accurately. These cases of failed incubation temperature data collection were identified by visual inspection of the incubation temperature time series blind to factors in the analysis and were removed from the dataset. When this occurred, we discarded the affected days of observation until the following iButton exchange. Our incubation analyses only included days of incubation after the clutch was completed and started no earlier than 15 days before the hatch date. From a total of 1,283 days of observations, a maximum of 729 days of incubation temperature recordings from 102 clutches were included in the analysis (sample sizes vary slightly across statistical models; details are given in the result section and Table S1).

Incubation behaviour (e.g., on-and off-bout timing) was determined using the R package incR (v1.1.0; Capilla-Lasheras 2018; Gwinner et al. 2018), choosing parameter values for incRscan validated for great tit incubation (Capilla-Lasheras 2018; lower.time=22, upper.time=3, sensitivity=0.15, temp.diff=8, maxNightVar_accepted=2). In short, to determine incubation on-and off-bouts, incRscan used variation in incubation temperatures during a time window (10pm to 3am, lower.time and upper.time parameters) in which females were assumed to incubate constantly, unless incubation temperatures dropped more than two degrees (maxNightVar_accepted parameter; see more details in Capilla-Lasheras 2018). For each incubating female, we determined: first morning off-bout, last evening on-bout, and duration of active day (e.g., time difference between the first morning off-bout and the last evening on-bout).

## Data analysis

### General modelling procedures

All analyses and visualisations were performed in R (version 4.2.1; R Core Team 2022). Generalised linear mixed models (GLMM) were employed to explain variation in several incubation and reproductive traits (see below). For each of these traits, we built a full model that contained all explanatory variables and interactions of interest for each trait (see below). Then, we used likelihood-ratio tests (LRTs) to assess the statistical importance of each model predictor. We removed non-significant interactions from the initial full models to ease biological interpretation of single effect predictors. However, we did not apply model simplification beyond non-significant interactions and present the resulting full model outputs. Linear and quadratic terms were retained in all models and fitted using orthogonal polynomials to improve model convergence and assess their statistical importance independently. Random effects were present in every model as specified for the analysis of each response variable (details below). We formally tested for non-zero model slopes of interactive terms using Wald χ^2^ tests implemented in the R package car (v3.1.0; Fox & Weisberg, 2019) via its linearHypothesis function. All statistical models were performed using the R package lme4 (v1.1.29; Bates et al. 2015). Gaussian model residuals were visually inspected to check the assumption of normality using the R package performance (v0.10.1; Lüdecke et al., 2021). The R package DHARMa (v0.4.5; Hartig 2018) was employed to check the normality of residuals in non-gaussian models.

### Statistical models

Incubation behaviour: We analysed clock (i.e., time after midnight) and relative (i.e., time relative to sunrise or sunset time) onset and end of diel activity. To account for differences in photoperiod throughout the breeding season, we calculated relative onset as the time of the first incubation off-bout minus sunrise time for each day (i.e., positive values represent onset of activity after sunrise, whereas negative vales indicate an onset of activity earlier than sunrise). Similarly, relative end of activity was defined per day as the time of the last on-bout minus sunset time (i.e., positive values represent end of activity after sunset, whereas negative values indicate an end of activity earlier than sunset). Full models for onset and end of activity (both clock and relative metrics) included as explanatory variables habitat (urban versus forest), clutch size (as a continuous predictor), mean daily temperature (as a continuous predictor), and days before hatching (as a continuous predictor whose minimum value was one [i.e., one day before hatching], included as a quadratic and a linear - these terms effectively modelled within-female variation in onset and end of activity). Additionally, we controlled for among-nest differences in timing of reproduction by including the date of incubation start (i.e., clutch completion date) as a fixed effect predictor (in number of days after April 1; included as a quadratic and a linear term - these terms effectively modelled among-female [e.g., cross-sectional] variation in onset and end of activity). Temporal predictors of incubation behaviour were included in the analysis as linear and quadratic terms given the evidence for negative quadratic temporal effects on incubation reported before (Cooper & Voss, 2013; Gwinner et al., 2018). We also included the interactions between habitat and days before hatching (both quadratic and linear terms), and between habitat and incubation start date (both quadratic and linear terms). Breeding attempt identity (included as a 90-level factor for nest-box identity, 79 out of the 90 nest-boxes included in the analysis [i.e., 87%] were used in a single year only), site (5-level factor) and year (3-level factor) were included as random effect intercepts. Using the same model structure, we analysed the duration of the active day of incubating females, defined as the time interval between the first incubation off-bout and the last on-bout per day.

We use the amount of variation explained by breeding attempt identity to calculate within-breeding-attempt consistency (e.g., repeatability) in female chronotype, but we do acknowledge that this calculation could be improved by tracking individual females across multiple breeding years (see Discussion). Specifically, consistency in female onset, end and duration of activity, was calculated as the proportion of variation explained by the breeding attempt identity random effect in the linear mixed models presented above (i.e., including year and site as random effects), as implemented in the R package rptR (Stoffel et al., 2017). Female chronotype for subsequent analyses (see below) was defined as the average within-nest onset of activity, but we also report consistency (e.g., repeatability) for end and duration of activity. We additionally analysed incubation start dates using a Gaussian GLMM with clutch size and habitat as fixed effects, and breeding attempt identity, site and year as random effects.

Survival of nestlings to fledging and nestling weight: A Poisson GLMM was used to explain variation in the number of nestlings that survived to fledging. The probability of total brood failure (i.e., the probability that no nestling survived to fledging) was modelled using a binomial GLMM. Given the lack of zero values (that Poisson distributions do have), an LMM was employed to analyse the number of nestlings that survived to fledging excluding broods in which no nestlings survived. Variation in the average 13-day old nestling weight per brood was analysed using an LMM. These models included habitat (urban versus forest), female chronotype (see definition above), hatching date (as a continuous variable in days after January 1; included in the model as a linear and a quadratic term) and clutch size as fixed effect predictors. The interactions between hatching date and habitat, and between female chronotype and habitat, were also added. Breeding attempt identity (90-level factor for survival analysis and 53-level factor in nestling weight analysis), site (5-level factor) and year (3-level factor) were included as random effect intercept.

### Ethical note

Nestlings were captured and minimally disturbed (for weighing) in their nest-boxes under ringing licenses granted to the authors by the British Trust of Ornithology. We adhered to the ASAB/ABS Guidelines for the use of animals in research. This project did not involve harmful manipulations of the study individuals or their environmental conditions.

## Results

### Correlates and consistency of incubating female chronotype

We recorded nest temperatures in 2016, 2017 and 2018, and analysed a maximum of 729 days of great tit incubation in 102 clutches (median = 7 days of incubation per clutch; range = 1 - 15 days; see details of sample sizes in Table S1). Urban great tit females laid their eggs and started incubation earlier in the year than forest females, and thus experienced shorter days with later sunrise and earlier sunset (start of incubation date: mean_urban_ ± SE = 30^th^ April ± 1.09 days, SD_urban_ = 1.09 days, N_urban_ = 27 clutches; mean_forest_ ± SE = 8^th^ May ± 0.61 days, SD_forest_ = 0.61 days, N_forest_ = 75 clutches; χ^2^ = 10.59, p = 0.001). Therefore, we detail results separately for the two habitats, but all data were analysed together in overarching models.

We found that, at the population level, clock time of activity onset was affected by habitat and by the date when incubation started (interaction ‘Incubation start date × habitat’: χ^2^ = 6.02, p = 0.014; Table 1; Figure 1a & 1b). Urban females closely tracked the seasonally advancing sunrise time (Figure 1a), but forest females largely ignored this advance and started their activity at a similar time throughout the season (Figure 1b; slope in Figure 1b was not significantly different from zero, χ^2^ = 1.01, p = 0.315). Whereas early-breeding urban females started activity later than forest females, for late-breeding birds the pattern reversed, so that urban females started their day earlier than forest females (Figure 1a & 1b). In contrast, at the population level, end of activity was similar in both habitats and became progressively earlier with later incubation start date (Figure S2; Table S2). Overall, the active day lengthened over the breeding season for urban but tended to shorten for forest females (Figure S3, Table S4). These patterns at the population level for clock time of onset and end of activity were broadly matched by temporal variation within clutches (i.e., variation between the first and last day of incubation of a clutch; Tables 1, S2, Figures S4, S5). Ambient temperature and clutch size did not affect clock time of onset and end of activity (Tables 1 and S2).

**Figure 1.**
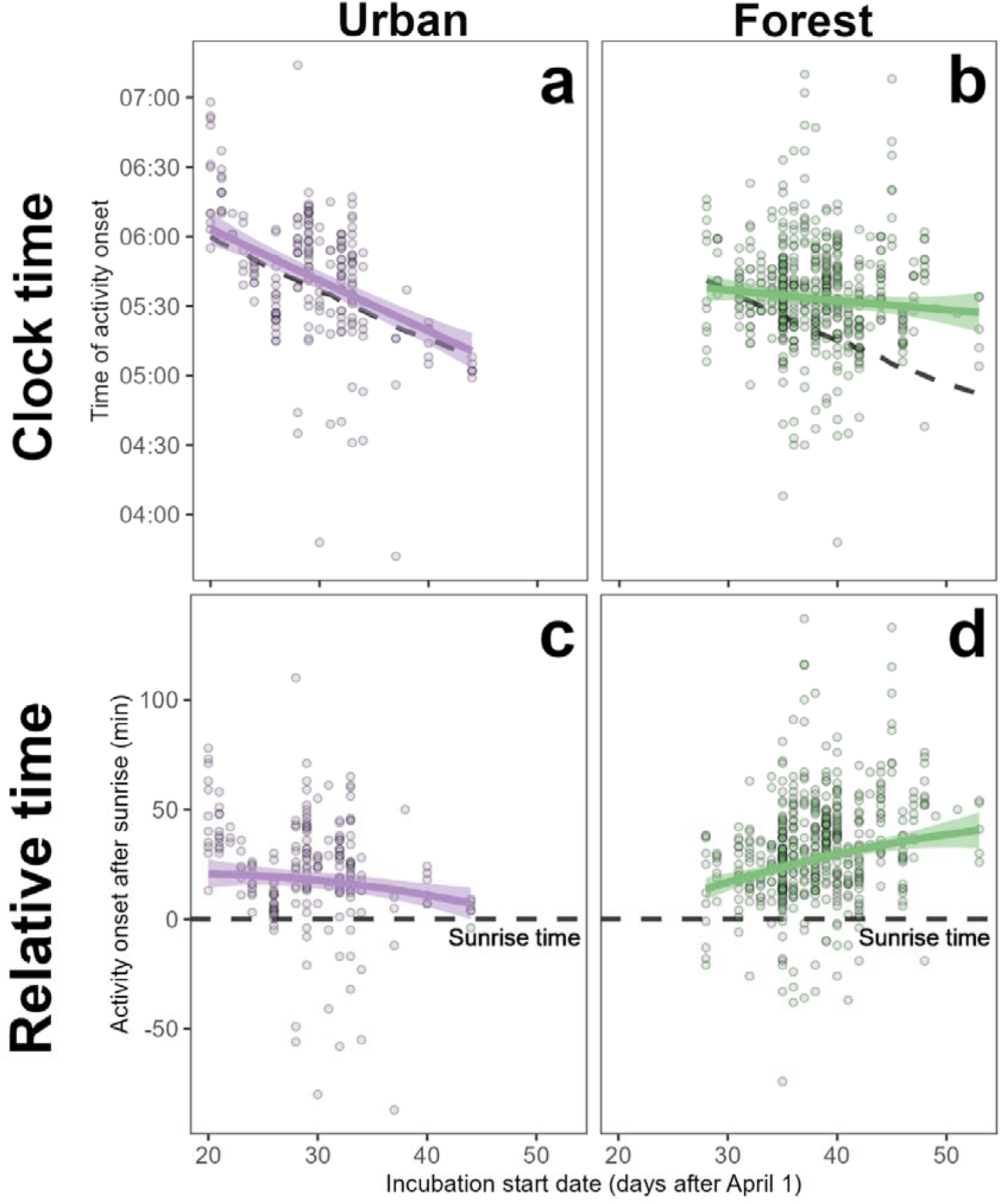
Clock and relative time of activity onset in urban and forest great tit females. **(a)** At the population level, urban females advanced their clock time of activity onset throughout the breeding season, seemingly tracking the seasonal advance of sunrise time (dashed line); whereas **(b)** the onset of activity of forest females remained relatively constant throughout the breeding season. Consequently, **(c)** urban females modified their onset of activity relative to sunrise over the breeding season only slightly, while forest females started their activity progressively later relative to sunrise. Points represent raw data while thick solid lines and shaded areas provide mean model predictions ± 1 SE (see model coefficients in Tables 1 and 2).

**Table 2.**
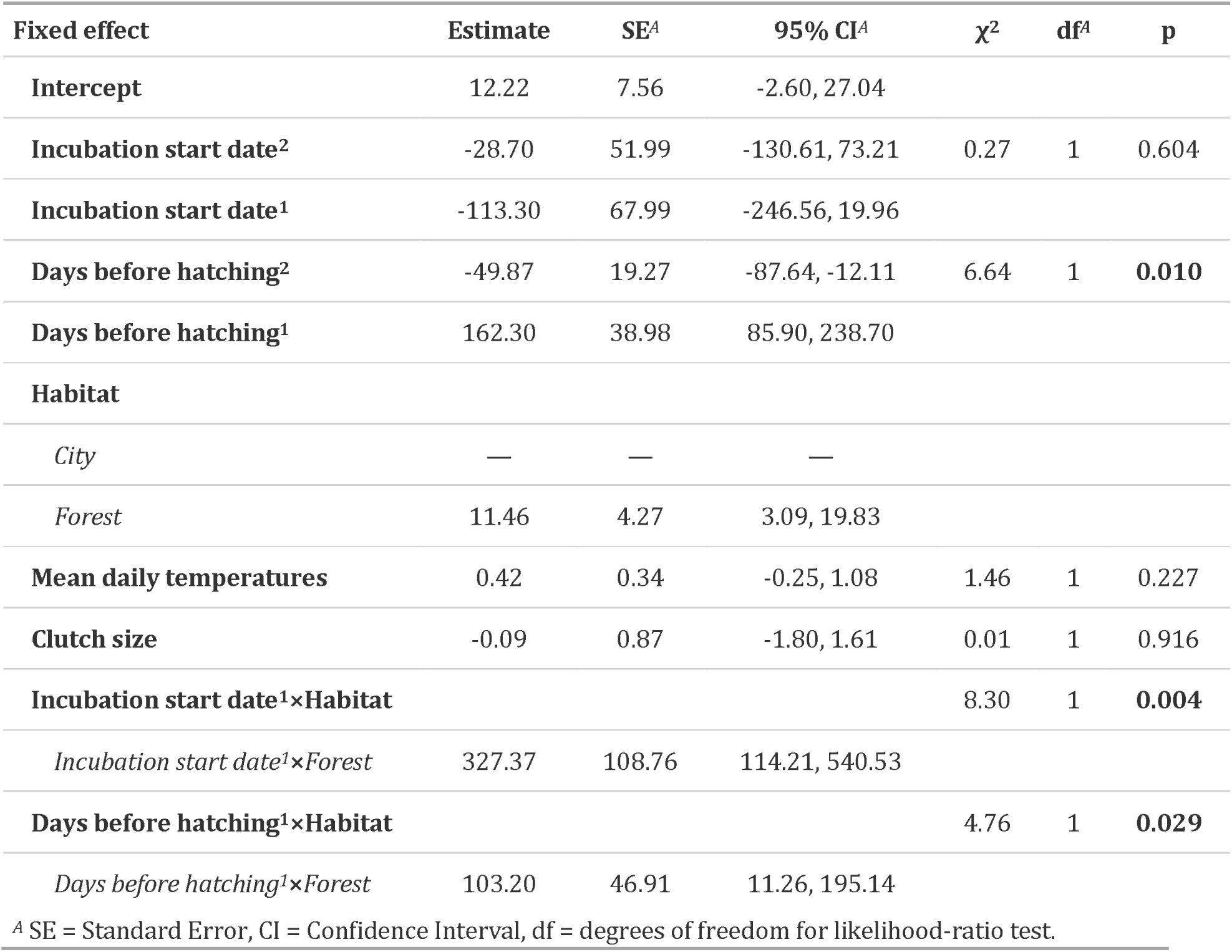
Likelihood-ratio test (LRT) results and model coefficients for predictors explaining variation in relative time of female onset of activity (i.e., time of first incubation off-bout minus sunrise time; n = 729 days of incubation). Significant terms are highlighted in bold. LRT results for ‘Habitat’, ‘Incubation start Date^1^’ and ‘Days before hatching^1^’ are not provided as these terms were part of a significant interaction present in the final model. The interactions ‘Incubation start date^2^×Habitat’ (χ^2^ = 1.84, p = 0.175) and ‘Days before hatching^2^×Habitat’ (χ^2^ df = 1 = 0.68, p = 0.409) were not df = 1 significant and were dropped from the model. Model coefficients (‘Estimate’) for relative onset of activity (given in min after sunrise) are shown along with standard errors (‘SE’) and 95% confidence intervals. Superscripts ‘^1^’ and ‘^2^’ refer to linear and quadratic terms, respectively.

| Fixed effect | Estimate | SE<sup>A</sup> | 95% CI<sup>A</sup> | $\chi^2$ | df<sup>A</sup> | p |
| --- | --- | --- | --- | --- | --- | --- |
| **Intercept** | 12.22 | 7.56 | -2.60, 27.04 |  |  |  |
| **Incubation start date<sup>2</sup>** | -28.70 | 51.99 | -130.61, 73.21 | 0.27 | 1 | 0.604 |
| **Incubation start date<sup>1</sup>** | -113.30 | 67.99 | -246.56, 19.96 |  |  |  |
| **Days before hatching<sup>2</sup>** | -49.87 | 19.27 | -87.64, -12.11 | 6.64 | 1 | **0.010** |
| **Days before hatching<sup>1</sup>** | 162.30 | 38.98 | 85.90, 238.70 |  |  |  |
| **Habitat** |  |  |  |  |  |  |
| *City* | — | — | — |  |  |  |
| *Forest* | 11.46 | 4.27 | 3.09, 19.83 |  |  |  |
| **Mean daily temperatures** | 0.42 | 0.34 | -0.25, 1.08 | 1.46 | 1 | 0.227 |
| **Clutch size** | -0.09 | 0.87 | -1.80, 1.61 | 0.01 | 1 | 0.916 |
| **Incubation start date<sup>1</sup>×Habitat** |  |  |  | 8.30 | 1 | **0.004** |
| *Incubation start date<sup>1</sup>×Forest* | 327.37 | 108.76 | 114.21, 540.53 |  |  |  |
| **Days before hatching<sup>1</sup>×Habitat** |  |  |  | 4.76 | 1 | **0.029** |
| *Days before hatching<sup>1</sup>×Forest* | 103.20 | 46.91 | 11.26, 195.14 |  |  |  |

<sup>A</sup> SE = Standard Error, CI = Confidence Interval, df = degrees of freedom for likelihood-ratio test.

**Table 3.** Likelihood-ratio test (LRT) results and model coefficients for relative female chronotype and other predictors hypothesised to explain variation in nestling survival to fledging (n = 101 broods). The interactions ‘Days before hatching^1^×Habitat’ (χ^2^ = 0.05, p = 0.831), ‘Days before hatching^2^×Habitat’ (χ^2^ df = 1 = 0.01, p = 0.927) and ‘Female chronotype×Habitat’ (χ^2^ df = 1 = 0.48, p = 0.489) were not significant and were dropped df = 1 from the model. Model coefficients (‘Estimate’) are shown in their link scale (logit) along with standard errors (‘SE’) and 95% confidence intervals (‘95% CI’). Superscripts ‘^1^’ and ‘^2^’ refer to linear and quadratic terms.

| Fixed effect | Estimate | SE<sup>A</sup> | 95% CI<sup>A</sup> | $\chi^2$ | df<sup>A</sup> | p |
| --- | --- | --- | --- | --- | --- | --- |
| **Intercept** | 0.14 | 0.27 | -0.38, 0.66 |  |  |  |
| **Hatching date<sup>1</sup>** | -0.08 | 0.82 | -1.69, 1.53 | 0.01 | 1 | 0.919 |
| **Hatching date<sup>2</sup>** | -0.37 | 0.59 | -1.53, 0.79 | 0.40 | 1 | 0.527 |
| **Female chronotype** | -0.11 | 0.05 | -0.22, 0.00 | 4.02 | 1 | **0.045** |
| **Habitat** |  |  |  | 5.12 | 1 | **0.024** |
| *City* | — | — | — |  |  |  |
| *Forest* | 0.38 | 0.17 | 0.05, 0.71 |  |  |  |
| **Clutch size** | 0.14 | 0.03 | 0.08, 0.21 | 18.66 | 1 | **<0.001** |

<sup>A</sup> SE = Standard Error, CI = Confidence Interval, df = degrees of freedom for likelihood-ratio test.

**Table 4.** Likelihood-ratio test (LRT) results and model coefficients for relative female chronotype and other predictors hypothesised to explain variation in pre-fledging weight of nestlings on day 13 (n = 57 broods). The interactions ‘Days before hatching^2^×Habitat’ (χ^2^ = 0.04, p = 0.847) and ‘Female chronotype×Habitat’ (χ^2^ = 0.02, p = 0.892) were not significant and were dropped from the model. Model coefficients (‘Estimate’) are shown along with standard errors (‘SE’) and 95% confidence intervals (‘95% CI’). Superscripts ‘^1^’ and ‘^2^’ refer to linear and quadratic terms.

| Fixed effect | Estimate | SE<sup>A</sup> | 95% CI<sup>A</sup> | $\chi^2$ | df<sup>A</sup> | p |
| --- | --- | --- | --- | --- | --- | --- |
| **Intercept** | 16.95 | 0.70 | 15.58, 18.32 |  |  |  |
| **Hatching date<sup>1</sup>** | -16.18 | 3.56 | -23.16, -9.20 |  |  |  |
| **Hatching date<sup>2</sup>** | -10.29 | 1.91 | -14.03, -6.55 | 19.88 | 1 | **<0.001** |
| **Female chronotype** | 0.06 | 0.14 | -0.21, 0.34 | 0.20 | 1 | 0.656 |
| **Habitat** |  |  |  |  |  |  |
| *City* | — | — | — |  |  |  |
| *Forest* | 2.69 | 0.52 | 1.67, 3.70 |  |  |  |
| **Clutch size** | -0.16 | 0.08 | -0.31, -0.01 | 4.05 | 1 | **0.044** |
| **Hatching date<sup>1</sup>×Habitat** |  |  |  | 10.87 | 1 | **0.001** |
| *Hatching date<sup>1</sup>×Forest* | 20.86 | 5.23 | 10.61, 31.11 |  |  |  |

<sup>A</sup> SE = Standard Error, CI = Confidence Interval, df = degrees of freedom for likelihood-ratio test.

Relative onset of activity depended on habitat and on the date when incubation started (Figure 1c-1d; Table 2). Females that initiated incubation later in the year began their day relative to sunrise progressively earlier in the city, but progressively later in the forest (interaction ‘Incubation start date × habitat’: χ^2^ = 8.30, p = 0.004; Figure 1c-1d). The end of activity relative to sunset advanced consistently over the season in both habitats (‘Incubation start date^1^’: χ^2^ = 30.44, p <0.001, ‘Incubation start date^2^’: χ^2^ = 0.20 p = 0.658; Figure S2c-S2d; Table S3). Ambient temperature and clutch size did not affect relative time of onset and end of activity (Tables 2, S3). These patterns at the population level for relative time of onset and end of activity were broadly matched by temporal variation within clutches (i.e., variation between the first and last day of incubation of a clutch; Tables 2, S3; Figures S4, S5).

We identified consistent individual differences in the time of onset of activity (i.e., female chronotype). Among-female differences explained 31 % of the variation in clock onset time (LRT on the breeding attempt ID random effect: χ^2^ = 133.18, p < 0.01 ; repeatability [95%CI] = 0.31 [0.21, 0.41]). Analyses of relative onset time (i.e., correcting for changes in sunrise time) yielded similar results, with consistent among-female differences in onset of activity (repeatability [95%CI] = 0.32 [0.22, 0.41]). We also found consistent among-female differences in the end of activity, both in clock (repeatability [95%CI] = 0.26 [0.17, 0.35]) and relative end of activity (repeatability [95%CI] = 0.27 [0.17, 0.35]); and consistent among-female differences in the duration of the active day (repeatability [95%CI] = 0.20 [0.13, 0.28]).

### Effects of female chronotype on fledging success and pre-fledging weight

We detected substantial variation between broods in the number of nestlings that survived to fledging. Relative female chronotype predicted the number of surviving nestlings: the earlier the female chronotype, the more nestlings fledged (χ^2^ = 4.02, p = 0.045; Figure 2a; Table 3). This effect was consistent across habitats (interaction between female chronotype and habitat, χ^2^ = 0.48, p = 0.489) and was robust to controlling for clutch size (χ^2^ = 18.66, p < 0.0001; Table 3). The number of surviving chicks was also affected by habitat (χ^2^ = 5.12, p = 0.024; Table 3): urban females fledged 0.54 nestlings less than forest females (i.e., a decrease in surviving nestlings of 46%; Table 3). Conversely, clock chronotype of females did not predict the number of surviving nestlings (Table S5).

**Figure 2.**
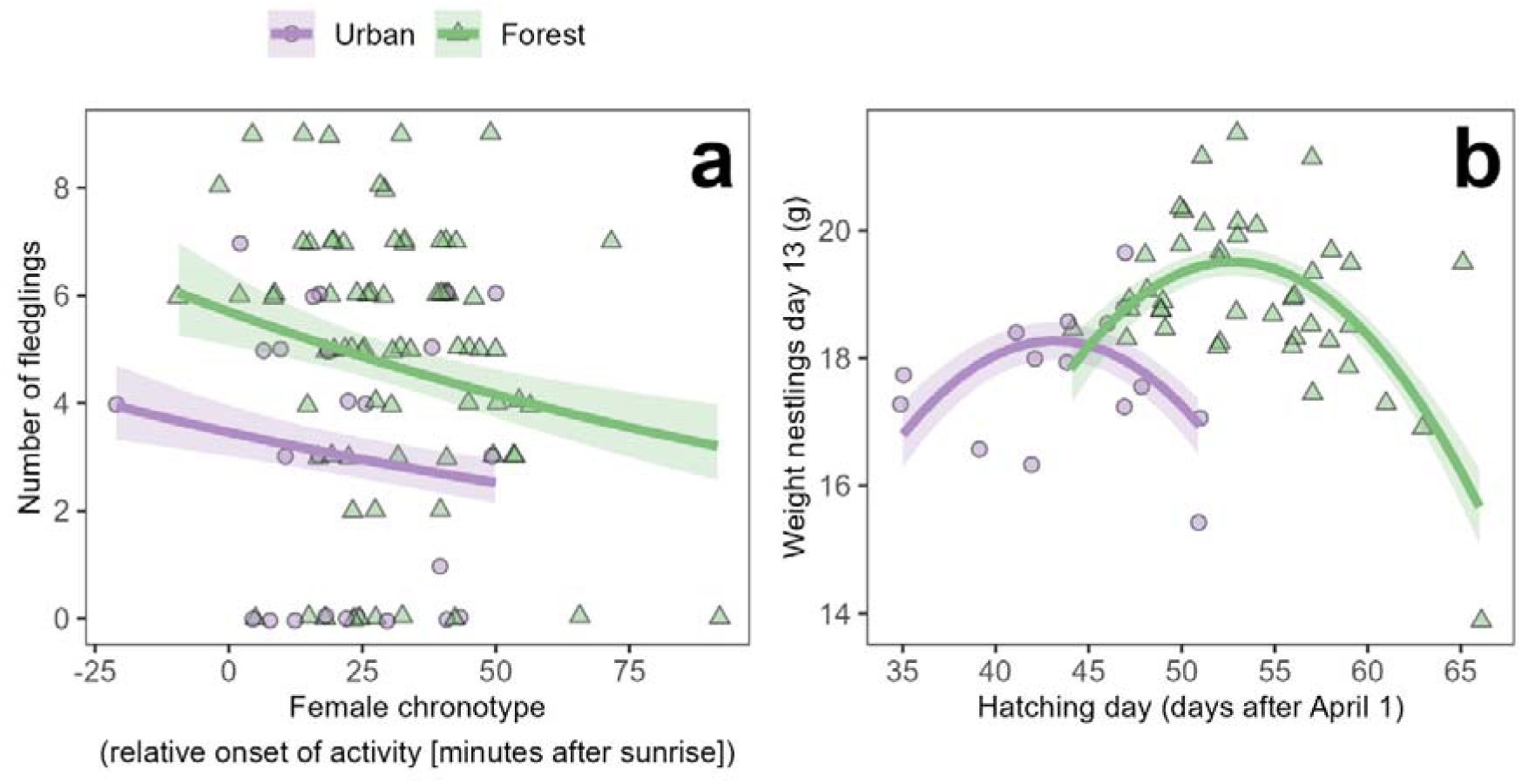
Effects of relative female chronotype and seasonal time on pre-fledging survival and weight. **(a)** Both in urban and forest nests, progressively later chronotype of the breeding female was negatively associated with the number of nestlings that survived to fledging (N = 101 broods; Table 3). **(b)** Urban nestlings on day 13 of their life were lighter than forest nestlings (N = 57 broods; Table 4). Variation in the weight of 13-day old nestlings showed a similar seasonal pattern in forest and city habitats. Points represent raw data, while thick lines and shaded areas provide mean model predictions ± 1 SE (Table 3 and 4).

To investigate the breeding parameters that could have generated the decreasing reproductive success with later relative female chronotype, we performed two additional analyses. Firstly, we assessed whether relative female chronotype was associated with total brood failure, and found no support (effect of relative female chronotype on the probability that no nestling survives to fledging: χ^2^ = 0.94, p = 0.333; Table S6). Secondly, we tested whether relative female chronotype predicted the number of nestlings surviving to fledging in successful broods (i.e., those that fledged at least one offspring), and confirmed that earlier relative chronotypes fledged more offspring than later chronotypes (χ^2^ = 4.45, p = 0.035; Table S7).

Mean body mass of 13-day old nestlings was affected by habitat, hatching date and clutch size. Forest nestlings were on average 2.69 g (95%CI = [1.67, 3.70]; Table 4) heavier than urban nestlings of the same age. In both habitats, pre-fledging weight was higher for broods that hatched in the middle of the season (Figure 2b), and nestlings from larger clutches were on average lighter (χ^2^ = 4.05, p = 0.044; Table 4). Neither relative nor clock female chronotype affected pre-fledging weight in either habitat (relative chronotype: χ^2^ = 0.20, p = 0.656; clock chronotype: χ^2^ = 0.06, p = 0.801; interaction terms between chronotype and habitat were non-significant, for relative chronotype: χ^2^ = 0.48, p = 0.489; for clock chronotype: χ^2^ = 0.36, p = 0.551).

## Discussion

Recent research has identified surprisingly high variation in chronotype of free-living animals, but determinants and effects of this variation are still largely unclear. Our study is among the few that have identified fitness correlates of (relative) chronotype in female animals. We firstly show high repeatability of timing, and thus corroborate evidence of chronotype as a consistent individual trait in birds, including in our study species (Graham et al., 2017; Meijdam et al., 2022; Stuber et al., 2015). We then show that the relative chronotype of female great tits, measured during the incubation period, predicted reproductive success, such that early-rising females raised more offspring to fledging than late (relative) chronotypes. As expected based on previous studies (Capilla-Lasheras et al., 2022; Dominoni et al., 2013), we have also found that urban great tits breed earlier in the season than non-urban great tits.

Early rising may be beneficial to replenish energy stores after night-time, and the ability of small passerine birds to successfully forage peaks in the early morning, once light conditions are suitable (Kacelnik, 1979; Pagani-Núñez & Senar, 2016). However, foraging in the early morning can be costly because of low ambient temperatures and increased predation risk (Mcnamara et al., 1994). It has been speculated that activity onset may be influenced by the condition of an individual, in addition to its endogenous circadian clock. However, the direction of such an influence is still unclear. For example, early song production has been interpreted as an honest signal of male quality, suggesting that superior condition is required for an early start of the day (Murphy et al., 2008). Conversely, in incubating females, boosting energy reserves through warming of the nest led to longer night rest (Arct et al., 2022; Bryan & Bryant, 1999; Gwinner et al., 2018). Likely, possible links between a bird’s condition and rising time are sensitive to ecological context.

Balancing costs and benefits of early rising might be intricate during incubation. For uniparental incubators, self-maintenance is weighed up against maximal offspring development (Nord & Cooper, 2020). This trade-off is heightened during early morning hours when incubators must replenish energy stores. Yet, because the typically low morning temperatures risk cooling of the eggs, an incubating female should delay leaving the nest until she can forage efficiently. Early rising may thus indicate superior foraging skills of incubating females, as proposed for courtship song and provisioning of males (McNamara et al., 1987; Murphy et al., 2008; Pagani-Núñez & Senar, 2016). If so, the higher reproductive success we found for early-rising females might be an indirect result of the females’ superior condition, as previously proposed for early-breeding females (Verhulst and Nilsson 2008) (A. Phillimore, pers. observation) Despite the fact that our results do not support this idea, it is also possible that early rising might be indicative of females in poor condition that cannot tolerate further depletion of energy and, hence, need to leave the nest early when eggs are at high risk of cooling (Nord & Cooper, 2020).

The ability to perform efficiently early in the day likely also depends on circadian mechanisms that facilitate an early start, as demonstrated in human athletes (Vitale & Weydahl, 2017). Reproductive advantages due to circadian-based early-rising have been proposed for male great tits whose circadian rhythm affects extra-pair paternity (EPP) and have been supported by experiments on the same study species (Greives et al., 2015; Hau et al., 2017; Helm & Visser, 2010). Great tit chicks with fast circadian clocks were significantly more likely to be sired through EPP, and males whose circadian system was pharmacologically delayed lost paternity (Greives et al., 2015; Helm & Visser, 2010). As in these other studies, our work found benefits for the early bird, without indicating what benefits or costs, in turn, might arise for late chronotypes. A putative circadian basis to early chronotype could involve several mechanistic features. These include a fast clock (i.e., short free-running period; Helm & Visser 2010), but also individual variation in sensitivity to light (Brown et al., 2008; Helm et al., 2017; Jones et al., 2019; Tudorache et al., 2018). A contribution of light response pathways to the chronotype - fitness link is suggested by our findings for clock and relative timing. Fitness effects were evident only for chronotype relative to sunrise, whereas the clock time of activity onset showed no association.

We detected unexpected differences in response to sunrise, but not sunset, between females at urban and forest sites. Forest females started activity at almost constant times of day, despite the rapid advance of sunrise time over the breeding season. Conversely, urban females were far more responsive to light and largely tracked the rapid advance of sunrise. This finding was counter to the expectation that in urban habitats, where artificial light at night is prevalent, the birds’ responsiveness to natural light changes would be reduced (Dominoni et al., 2013; Roenneberg et al., 2007), or that, like some species under continuous light, birds in light-polluted areas might not use light conditions to time their activities (Huffeldt & Merkel, 2016). It is possible that habitat differences other than light levels contributed to the differences in behaviour. For related blue tits (Cyanistes caeruleus), the same study habitats differed in quality, with poorer adult state and reproductive success in the city (Capilla-Lasheras et al., 2017; Pollock et al., 2017). Thus, some urban great tit females may have needed to forage at the earliest opportunity to replenish their resources, without an apparent impact on reproductive success. Disentangling effects of the internal circadian clock on chronotype from those of the birds’ body condition would require experimental examination (Dominoni et al., 2013; Greives et al., 2015).

The only other study we are aware of that examined reproductive success relative to incubation chronotype did not find such an association (Maury et al., 2020). This investigation differed in several aspects, including use of the European starling (Sturnus vulgaris) as a study species. While we cannot explain the different findings, we speculate that colonial breeding of the studied starlings may have affected synchronicity, and thereby altered or obscured effects of chronotype (Gwinner, 1966; Menaker & Eskin, 1966). In other contexts, fitness implications of chronotype are also beginning to arise. For example, a recent study on fish showed that under fishery pressure, chronotype was associated with differential survival (Martorell-Barceló et al., 2018). Still, we are far from understanding how variation in chronotype is maintained.

Our study results come with some caveats. Because we report correlative data from wild birds, we cannot assess whether chronotype was affected by the local micro-environment, either directly or via differences in individual quality (Diez-Méndez, Cooper, et al., 2021; Maury et al., 2020). We have recorded female chronotype only during one life-cycle stage, incubation, similar to earlier studies on males that considered only courtship (Murphy et al., 2008). Thus, the consistency of chronotype across life stages remains to be tested. Similarly, we inferred chronotype from onset of activity across multiple days of the same breeding event, and we could not assess consistency of chronotype for the same female across multiple breeding seasons. Correlated environmental conditions or female body condition within breeding attempts could have potentially increased the estimate of chronotype consistency. Comparing our estimates of chronotype with quantifications from studies that track individuals across multiple breeding seasons will greatly expand the significance of our results and shed new light on the environmental contributions to chronotype.

Nonetheless, our study strengthens the evidence for variation in chronotype in free-living animals and provides a sought-after link to reproductive success. We extend the circadian focus of chronotype studies by indicative findings on light pathways, and confirm the importance of looking at both relative and clock time, as previously suggested for avian incubation (Shaw & Cresswell, 2014). Future challenges, likely requiring experimental approaches, are a disentangling of effects of endogenous clock from body condition, and determination of counter-balancing benefits that maintain variation in chronotype.

## Data availability

All R scripts and datasets needed to reproduce the analyses presented in this paper are available at: https://github.com/PabloCapilla/incubation_chronotype. Should the manuscript be accepted, a DOI to this data repository will be provided.

## Author contributions

RJW and BH conceived of the study. RJW, CLOM, DMD and BH collected the data. PC-L performed all statistical analyses, with input from RJW, CLOM, DMD and BH. PC-L and BH wrote the first draft of the manuscript, with major contributions by RJW. All authors read and revised the manuscript.

## Acknowledgements

We thank Ally Phillimore and an anonymous reviewer for their comments on a previous version of the manuscript.

