## Supplementary materials for "Reproductive fitness is associated with female chronotype in a songbird"

**Supplementary tables**

**Table S1**. Sample size in urban and forest habitats across the three years of study for each trait investigated.

| **Trait** | **year** | **Urban** | **Forest** | **Units** |
| --- | --- | --- | --- | --- |
| Onset of activity | 2016 | 14 (4) | 216 (30) | Days of observation (number of clutches) |
|  | 2017 | 103 (11) | 203 (33) | Days of observation (number of clutches) |
|  | 2018 | 89 (12) | 104 (12) | Days of observation (number of clutches) |
| End of activity | 2016 | 14 (4) | 209 (30) | Days of observation (number of clutches) |
|  | 2017 | 102 (11) | 201 (33) | Days of observation (number of clutches) |
|  | 2018 | 84 (12) | 97 (12) | Days of observation (number of clutches) |
| Duration of activity | 2016 | 14 (4) | 209 (30) | Days of observation (number of clutches) |
|  | 2017 | 102 (11) | 201 (33) | Days of observation (number of clutches) |
|  | 2018 | 84 (12) | 97 (12) | Days of observation (number of clutches) |
| Nestling survival | 2016 | 4 | 30 | number of clutches |
|  | 2017 | 11 | 33 | number of clutches |
|  | 2018 | 11 | 12 | number of clutches |
| Fledgling weight | 2016 | 4 | 24 | number of broods |
|  | 2017 | 6 | 10 | number of broods |
|  | 2018 | 5 | 8 | number of broods |

**Table S2.** Likelihood-ratio test (LRT) results and model coefficients for predictors explaining variation in female clock time end of activity (i.e., time of last daily incubation on-bout; n = 707 days of incubation). Significant terms are highlighted in bold. LRT results for ‘Habitat’ and ‘Days before hatching^1^’ are not provided as these terms were part of a significant interaction present in the model. The interactions ‘Incubation start date^2^×Habitat’ (χ^2^_df = 1_ = 0.60, p = 0.440), ‘Incubation start date^1^×Habitat’ (χ^2^_df = 1_ = 0.13, p = 0.719) and ‘Days before hatching^2^×Habitat’ (χ^2^_df = 1_ = 1.84, p = 0.175) were not significant and were dropped from the model. Model coefficients (‘Estimate’) for clock time end of activity (model coefficients are given in min after 00:00 h) are shown along with standard errors (‘SE’) and 95% confidence intervals (95% CI). Superscripts ‘1’ and ‘2’ refer to linear and quadratic terms, respectively.

| **Fixed effect** | **Estimate** | **SE***^A^* | **95% CI***^A^* | **χ^2^** | **df*^A^*** | **p** |
| --- | --- | --- | --- | --- | --- | --- |
| **Intercept** | 1,155.28 | 12.71 | 1,130.38, 1,180.18 |  |  |  |
| **Incubation start date^2^** | -32.84 | 62.95 | -156.21, 90.54 | 0.27 | 1 | 0.603 |
| **Incubation start date^1^** | -270.15 | 69.80 | -406.95, -133.34 | 9.08 | 1 | **0.003** |
| **Days before hatching^2^** | 19.59 | 34.98 | -48.97, 88.15 | 0.31 | 1 | 0.576 |
| **Days before hatching^1^** | 105.38 | 70.80 | -33.39, 244.15 |  |  |  |
| **Habitat** |  |  |  |  |  |  |
| *Urban* | — | — | — |  |  |  |
| *Forest* | 20.90 | 6.79 | 7.60, 34.20 |  |  |  |
| **Mean daily temperatures** | -0.07 | 0.62 | -1.28, 1.14 | 0.01 | 1 | 0.904 |
| **Clutch size** | -2.59 | 1.43 | -5.40, 0.23 | 3.23 | 1 | 0.072 |
| **Days before hatching^1^× Habitat** |  |  |  | 15.93 | 1 | **<0.001** |
| *Days before hatching^1^***×***Forest* | -342.66 | 85.19 | -509.63, -175.68 |  |  |  |
| *^A^* SE = Standard Error, CI = Confidence Interval, df = degrees of freedom for likelihood-ratio test. | | | | | | |

**Table S3.** Likelihood-ratio test (LRT) results and model coefficients for predictors explaining variation in female relative end of activity (i.e., time of last incubation on-bout minus sunset time; n = 707 days of incubation). Significant terms are highlighted in bold. LRT results for ‘Habitat’ and ‘Days before hatching^1^’ are not provided as these terms were part of a significant interaction present in the model. The interactions ‘Incubation start date^2^×Habitat’ (χ^2^_df = 1_ = 0.35, p = 0.557), ‘Incubation start date^1^×Habitat’ (χ^2^_df = 1_ = 0.35, p = 0.553) and ‘Days before hatching^2^×Habitat’ (χ^2^_df = 1_ = 1.96, p = 0.162) were not significant and were dropped from the model. Model coefficients (‘Estimate’) for relative time of end of activity (model coefficients are given in min after sunset) are shown along with standard errors (‘SE’) and 95% confidence intervals (95% CI). Superscripts ‘1’ and ‘2’ refer to linear and quadratic terms, respectively.

| **Fixed effect** | **Estimate** | **SE***^A^* | **95% CI***^A^* | **χ^2^** | **df*^A^*** | **p** |
| --- | --- | --- | --- | --- | --- | --- |
| **Intercept** | -118.84 | 12.90 | -144.12, -93.55 |  |  |  |
| **Incubation start date^2^** | -29.95 | 64.22 | -155.82, 95.93 | 0.20 | 1 | 0.658 |
| **Incubation start date^1^** | -520.10 | 73.64 | -664.44, -375.76 | 30.44 | 1 | **<0.001** |
| **Days before hatching^2^** | 21.25 | 35.15 | -47.64, 90.15 | 0.36 | 1 | 0.548 |
| **Days before hatching^1^** | -80.29 | 71.23 | -219.90, 59.32 |  |  |  |
| **Habitat** |  |  |  |  |  |  |
| *Urban* | — | — | — |  |  |  |
| *Forest* | 13.64 | 7.05 | -0.18, 27.46 |  |  |  |
| **Mean daily temperatures** | -0.10 | 0.62 | -1.32, 1.12 | 0.02 | 1 | 0.875 |
| **Clutch size** | -2.65 | 1.45 | -5.49, 0.18 | 3.33 | 1 | 0.068 |
| **Days before hatching^1^×Habitat** |  |  |  | 14.36 | 1 | **<0.001** |
| *Days before hatching^1^***×***Forest* | -328.15 | 85.67 | -496.07, -160.24 |  |  |  |
| *^A^* SE = Standard Error, CI = Confidence Interval, df = degrees of freedom for likelihood-ratio test. | | | | | | |

**Table S4**. Likelihood-ratio test (LRT) results and model coefficients for predictors explaining variation in the duration of the active day of incubating great tit females, calculated from clock time (i.e., time of last daily incubation on-bout minus time of first daily incubation off-bout; n = 707 days of incubation). Significant terms are highlighted in bold. LRT results for ‘Habitat’, ‘Incubation start date^1^’ and ‘Days before hatching^1^’ are not provided as these terms were part of a significant interaction present in the model. The interactions ‘Incubation start date^2^×Habitat’ (χ^2^_df = 1_ = 0.25, p = 0.616), ‘Incubation start date^1^×Habitat’ (χ^2^_df = 1_ = 1.979, p = 0.160) and ‘Days before hatching^2^×Habitat’ (χ^2^_df = 1_ = 0.756, p = 0.385) were not significant and were dropped from the model. Model coefficients (‘Estimate’) are shown along with standard errors (‘SE’) and 95% confidence intervals (95% CI). Superscripts ‘^1^’ and ‘^2^’ refer to linear and quadratic terms respectively.

| **Fixed effect** | **Estimate** | **SE***^A^* | **95% CI***^A^* | **χ^2^** | **df*^A^*** | **p** |
| --- | --- | --- | --- | --- | --- | --- |
| **Intercept** | 828.65 | 13.79 | 801.62, 855.68 |  |  |  |
| **Incubation start date^2^** | -144.54 | 66.08 | -274.05, -15.02 | 4.71 | 1 | **0.030** |
| **Incubation start date^1^** | -32.34 | 75.67 | -180.66, 115.97 | 0.18 | 1 | 0.670 |
| **Days before hatching^2^** | 67.10 | 40.34 | -11.97, 146.17 | 2.76 | 1 | 0.097 |
| **Days before hatching^1^** | 153.84 | 81.12 | -5.15, 312.84 |  |  |  |
| **Habitat** |  |  |  |  |  |  |
| *Urban* | — | — | — |  |  |  |
| *Forest* | 14.30 | 7.11 | 0.36, 28.24 |  |  |  |
| **Mean daily temperatures** | -0.60 | 0.71 | -1.99, 0.79 | 0.72 | 1 | 0.396 |
| **Clutch size** | -2.59 | 1.52 | -5.56, 0.39 | 2.90 | 1 | 0.088 |
| **Days before hatching^1^×Habitat** |  |  |  | 23.22 | 1 | **<0.001** |
| *Days before hatching^1^×Forest* | -473.36 | 97.43 | -664.32, -282.41 |  |  |  |
| *^A^* SE = Standard Error, CI = Confidence Interval, df = degrees of freedom for likelihood-ratio test. | | | | | | |

**Table S5**. Likelihood-ratio test (LRT) results and model coefficients for clock time female chronotype and other predictors hypothesised to explain variation in nestling survival to fledging (n = 101 broods). Model coefficients (‘Estimate’) are shown in their link scale (logit) along with standard errors (‘SE’) and 95% confidence intervals (‘95% CI’). Superscripts ‘^1^’ and ‘^2^’ refer to linear and quadratic terms.

| **Fixed effect** | **Estimate***^A^* | **SE***^A^* | **95% CI***^A^* | **χ^2^** | **df*^A^*** | **p** |
| --- | --- | --- | --- | --- | --- | --- |
| **Intercept** | 0.18 | 0.27 | -0.35, 0.70 |  |  |  |
| **Hatching date^1^** | -0.49 | 0.86 | -2.17, 1.20 | 0.32 | 1 | 0.572 |
| **Hatching date^2^** | -0.60 | 0.59 | -1.74, 0.55 | 1.07 | 1 | 0.300 |
| **Clock female chronotype** | -0.01 | 0.06 | -0.13, 0.10 | 0.06 | 1 | 0.801 |
| **Habitat** |  |  |  | 4.32 | 1 | **0.038** |
| *Urban* | — | — | — |  |  |  |
| *Forest* | 0.36 | 0.17 | 0.02, 0.69 |  |  |  |
| **Clutch size** | 0.14 | 0.03 | 0.08, 0.21 | 17.04 | 1 | **<0.001** |
| *^A^* SE = Standard Error, CI = Confidence Interval, df = degrees of freedom for likelihood-ratio test. | | | | | | |

**Table S6**. Likelihood-ratio test (LRT) results and model coefficients for predictors explaining variation in total brood failure (i.e., probability that no nestling survives to fledging; n = 101 broods). Superscripts ‘^1^’ and ‘^2^’ refer to linear and quadratic terms respectively.

| **Fixed effect** | **Estimate***^A^* | **SE***^A^* | **95% CI***^A^* | **χ^2^** | **df*^A^*** | **p** |
| --- | --- | --- | --- | --- | --- | --- |
| **Intercept** | -0.72 | 1.37 | -3.41, 1.97 |  |  | 0.598 |
| **Hatching date^1^** | -3.31 | 4.38 | -11.89, 5.26 | 0.58 | 1 | 0.447 |
| **Hatching date^2^** | -1.78 | 3.35 | -8.33, 4.78 | 0.30 | 1 | 0.587 |
| **Female chronotype** | 0.29 | 0.30 | -0.30, 0.88 | 0.94 | 1 | 0.333 |
| **Habitat** |  |  |  | 1.18 | 1 | 0.277 |
| *Urban* | — | — | — |  |  |  |
| *Forest* | -0.86 | 0.78 | -2.39, 0.67 |  |  |  |
| **Clutch size** | -0.05 | 0.19 | -0.41, 0.31 | 0.07 | 1 | 0.796 |
| *^A^* SE = Standard Error, CI = Confidence Interval, df = degrees of freedom for likelihood-ratio test. | | | | | | |

**Table S7.** Likelihood-ratio test (LRT) results and model coefficients for predictors explaining variation in the number of nestlings that survived to fledging, after excluding broods in which no nestling fledged (i.e., excluding total failure broods; n = 82 broods). Superscripts ‘^1^’ and ‘^2^’ refer to linear and quadratic terms respectively.

| **Fixed effect** | **Estimate** | **SE***^A^* | **95% CI***^A^* | **χ^2^** | **df*^A^*** | **p** |
| --- | --- | --- | --- | --- | --- | --- |
| **Intercept** | -0.41 | 0.73 | -1.85, 1.02 |  |  |  |
| **Hatching date^1^** | -3.07 | 1.67 | -6.34, 0.20 | 3.32 | 1 | 0.068 |
| **Hatching date^2^** | -1.08 | 1.41 | -3.85, 1.68 | 0.59 | 1 | 0.443 |
| **Female chronotype** | -0.34 | 0.16 | -0.65, -0.03 | 4.45 | 1 | **0.035** |
| **Habitat** |  |  |  | 3.67 | 1 | 0.055 |
| *Urban* | — | — | — |  |  |  |
| *Forest* | 0.89 | 0.46 | -0.01, 1.78 |  |  |  |
| **Clutch size** | 0.73 | 0.09 | 0.55, 0.92 | 45.81 | 1 | **<0.001** |
| *^A^* SE = Standard Error, CI = Confidence Interval, df = degrees of freedom for likelihood-ratio test. | | | | | | |

**Supplementary figures**

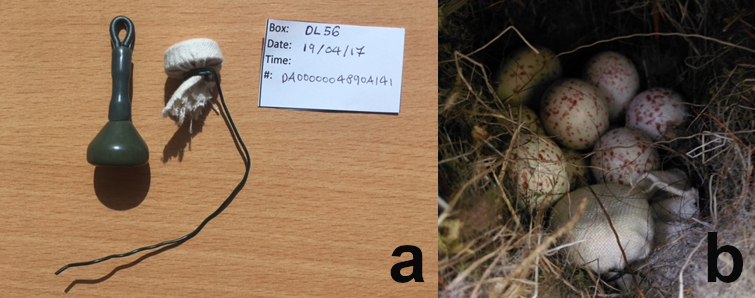

**Figure S1**. iButton setup for incubation data collection. (**a**) Pre-programmed iButton wrapped in fabric and attached to wire, 28.35g weight and iButton field label. (**b**) Detail of a nest cup with the iButton device positioned among great tit eggs.

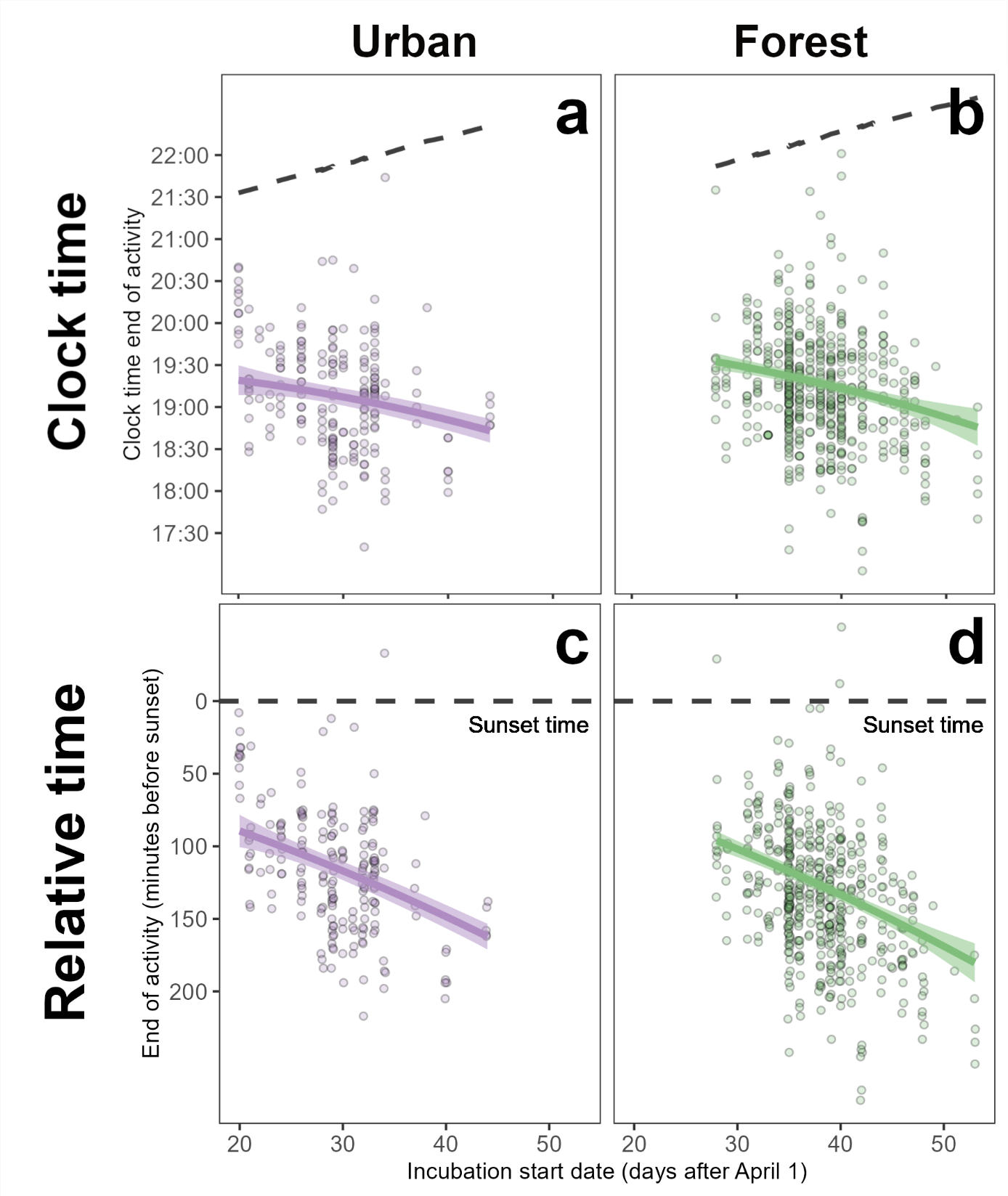

**Figure S2**. (**a**-**b**) Clock and (**c**-**d**) relative end of activity (i.e., time of the last incubation on-bout minus sunset time; positive values indicate minutes after sunset, whereas negative values indicate minutes before sunset) for incubating (**a**-**c**) urban and (**b**-**d**) non-urban female great tits throughout the breeding season. Points represent raw data, while thick solid lines and shaded areas provide mean model predictions ± 1 SE. Dashed line marks sunset time.

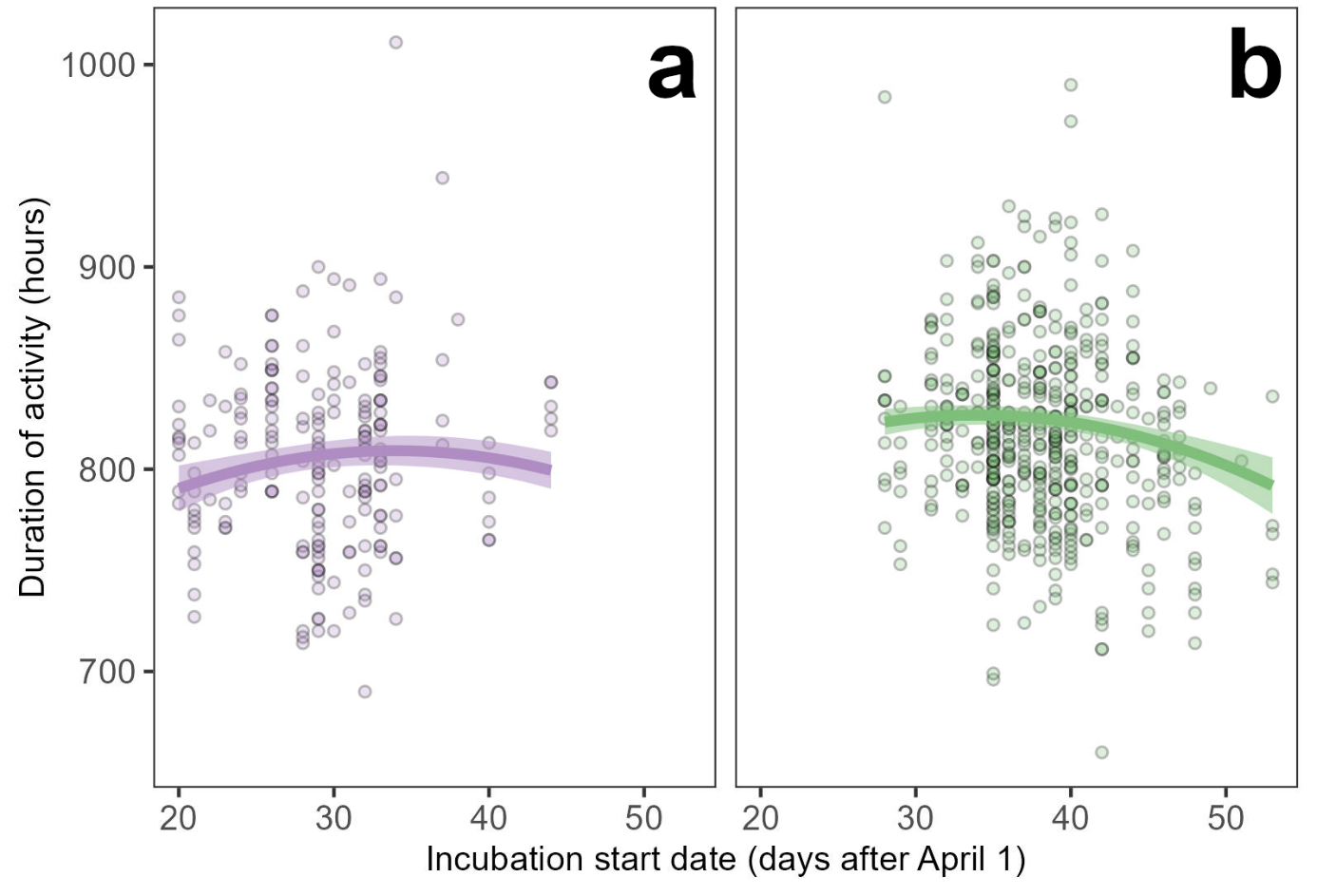

**Figure S3**. Duration of the active day (i.e., time difference between the first incubation off-bout and the last incubation on-bout per day) in incubating (**a**) urban and (**b**) non-urban female great tits throughout the breeding season. The active day lengthened over the breeding season for urban but tended to shorten for forest females as a consequence of differences in incubation start dates across habitats (i.e., this difference does not reflect a significant ‘Incubation start date × Habitat’ interaction; note that the shape of the quadratic fit is similar across habitats). Points represent raw data, while thick solid lines and shaded areas provide mean model predictions ± 1 SE (see model coefficients in Table S4).

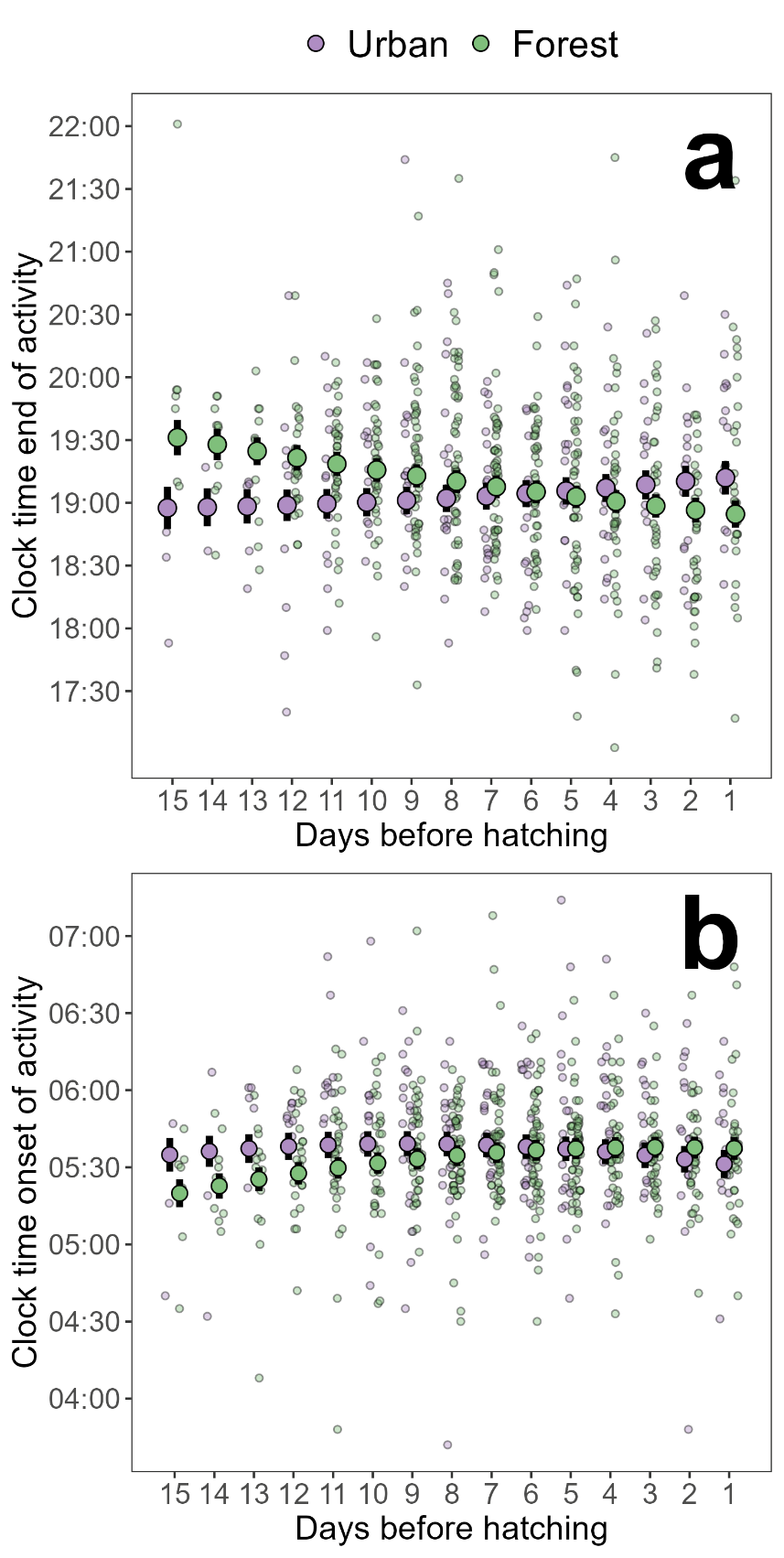

**Figure S4**. Clock end of activity (a) and onset of activity (b; i.e., time of the last incubation on-bout and time of the first incubation off-bout) per day of incubation (from 15 to 1 days before hatching; ‘0’ would indicate day of hatching) for Urban (in purple) and forest (in green) female great tits. Small transparent points represent raw data, while large solid points and bars provide mean model predictions ± 1 SE for each day of incubation (see model coefficients in Table 1 and Table S2). Dashed line marks sunset time.

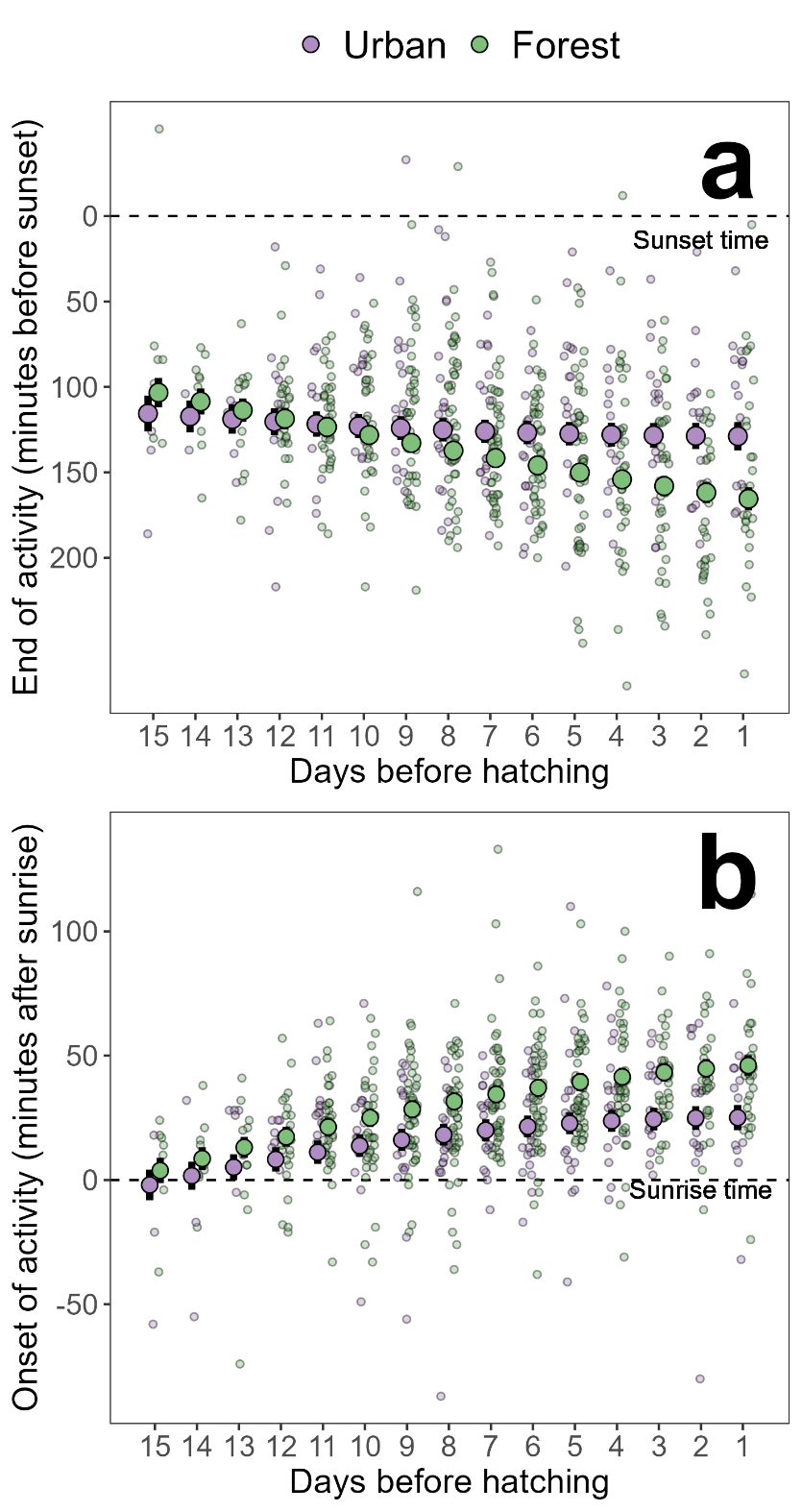

**Figure S5**. Relative end (a) and onset of activity (b; i.e., time of the last incubation on-bout minus sunset time, and time of the first incubation off-bout minus sunrise time; positive values indicate minutes after sunset/sunrise, whereas negative values indicate minutes before sunset/sunrise) per day of incubation (from 15 to 1 days before hatching; ‘0’ would indicate day of hatching) for urban (in purple) and forest (in green) female great tits. Small transparent points represent raw data, while large solid points and bars provide mean model predictions ± 1 SE for each day of incubation (see model coefficients in Table 1 and Table S3). Dashed line marks sunset time.
